# SPATIAL HABITAT DIFFERENCES DRIVE ABUNDANCE OF RED-CHEEKED CORDON-BLEU IN HUMAN-MODIFIED LANDSCAPES

**DOI:** 10.64898/2026.05.15.725372

**Authors:** Aminu Sani Khalifa

**Affiliations:** Department of Zoology, Faculty of Life Sciences, Federal University of Lafia, Nasarawa State, Nigeria

**Keywords:** *Uraeginthus bengalus*, habitat variation, farmland ecology, urban disturbance, savanna birds, bird abundance, Plateau State

## Abstract

Habitat modification is a major driver of avian population change in tropical savanna ecosystems. This study investigated habitat-related variation in the abundance of the Red-cheeked Cordon-bleu (*Uraeginthus bengalus*) across human settlements and surrounding farmlands in Laminga Village, Jos-East Local Government Area, Plateau State, Nigeria. Field surveys were conducted over a three-week period in November 2024 using 21 line transects sampled during peak bird activity periods. Bird abundance data were analysed using a Poisson Generalized Linear Model (GLM). Results showed that habitat type significantly influenced abundance, with significantly lower abundance recorded in human settlements compared to farmlands (β = −0.836, SE = 0.192, z = −4.359, p < 0.001). Transect length positively influenced abundance (β = 0.028, SE = 0.008, z = 3.600, p < 0.001). Model performance improved substantially from the null deviance (159.88) to the residual deviance (125.85), with an Akaike Information Criterion (AIC) value of 306.32. The findings suggest that farmlands provide more favourable habitat conditions for the species, likely due to greater vegetation availability and reduced structural disturbance relative to settlement areas. The study highlights the ecological importance of low-intensity agricultural landscapes in supporting avian persistence within human-modified savanna environments.

## INTRODUCTION

Bird populations are widely recognized as sensitive indicators of environmental change, particularly in landscapes increasingly modified by human activities, making them critical subjects in ecological monitoring and conservation research (Callaghan *et al*., 2024). Rapid global land-use transformation, driven by urban expansion, agricultural intensification, and infrastructure development, has significantly altered natural habitats, thereby influencing avian distribution, diversity, and abundance across different ecological zones (IPBES, 2021). Human-dominated landscapes, including settlements and farmlands, represent complex ecological systems where natural and artificial elements interact to shape species composition and ecological processes (Aronson *et al*., 2020; Shackleton *et al*., 2021). In sub-Saharan Africa, where human population growth and land conversion are accelerating, understanding how bird species respond to anthropogenic pressures is essential for biodiversity conservation and sustainable land management (Callaghan *et al*., 2024). Although several studies have examined bird responses to urbanization and agricultural practices globally, there remains a notable gap in localized studies focusing on specific species within rural African landscapes, particularly in Nigeria (Aronson *et al*., 2020). Laminga village in Jos-East Local Government Area represents a typical rural setting where human settlements coexist with agricultural lands, creating a mosaic of habitats with varying degrees of anthropogenic influence. Understanding how such environmental gradients affect the abundance of the Red-cheeked Cordon-bleu is essential for generating baseline ecological data and informing conservation strategies at the local level.

One of the defining features of human-dominated landscapes is habitat heterogeneity, which can both positively and negatively influence bird populations depending on species-specific ecological requirements (Morelli *et al*., 2021; Tryjanowski *et al*., 2020). Urban environments often contain a mosaic of built structures, green spaces, gardens, and remnant vegetation patches, creating novel ecosystems that differ significantly from natural habitats (Aronson *et al*., 2020; Shackleton *et al*., 2021). Some bird species, particularly generalists and synanthropic species, are able to exploit these environments by utilizing artificial structures for nesting and anthropogenic food sources for survival (Møller *et al*., 2021; Evans *et al*., 2022). In contrast, specialist species that depend on specific habitat conditions or resources may decline or disappear due to habitat loss and environmental disturbance (Benítez-López *et al*., 2021; Samia *et al*., 2022). Agricultural landscapes, which dominate large portions of rural areas, also play a crucial role in shaping avian ecology, particularly in developing regions where subsistence and small-scale farming are prevalent (Morelli *et al*., 2021). Farmlands can provide important foraging habitats for birds, especially granivorous and insectivorous species that benefit from crop residues, seeds, and associated invertebrate populations (Tryjanowski *et al*., 2020; Benton *et al*., 2021). However, agricultural intensification, including the use of agrochemicals, mechanization, and monoculture practices, has been linked to declines in bird abundance and diversity due to reduced habitat complexity and food availability (Tryjanowski *et al*., 2020). In less intensive systems, such as those commonly found in parts of sub-Saharan Africa, farmland heterogeneity may still support relatively diverse bird communities, although increasing pressure from land-use change poses ongoing risks (Evans *et al*., 2022).

The Red-cheeked Cordon-bleu (*Uraeginthus bengalus*) is a small estrildid finch widely distributed across sub-Saharan Africa, where it occupies a variety of open and semi-open habitats, making it a suitable model species for studying ecological responses to human-modified environments (BirdLife International, 2023). This species is particularly common in savanna regions, cultivated lands, village outskirts, and areas with scattered shrubs and grasses, reflecting its ecological flexibility and tolerance to moderate levels of disturbance (Craig *et al*., 2020). Its widespread occurrence across both natural and anthropogenic landscapes has made it an important subject in avian ecological studies focusing on habitat use, population dynamics, and adaptation to environmental change (Evans *et al*., 2022). The feeding ecology of the Red-cheeked Cordon-bleu is primarily centered on ground foraging, where individuals search for seeds in grasses and on bare soil surfaces, often in pairs or small flocks (Craig *et al*., 2020; Evans *et al*., 2022). In addition to seeds, the species may occasionally consume small insects, particularly during the breeding season when protein requirements increase for chick development (Morelli *et al*., 2021). This dietary flexibility enhances its ability to exploit a range of habitats, including those influenced by agricultural activities, where food resources may fluctuate seasonally due to planting and harvesting cycles (Tryjanowski *et al*., 2020). Habitat preference in *Uraeginthus bengalus* is strongly influenced by the availability of ground vegetation, seed resources, and suitable nesting sites, which are often found in heterogeneous landscapes combining natural and modified elements (Morelli *et al*., 2021; Redhead *et al*., 2021). Areas with moderate grass cover, scattered shrubs, and minimal disturbance tend to support higher densities of the species, while highly urbanized or heavily disturbed environments may reduce its abundance (Benítez-López *et al*., 2021). This pattern reflects the importance of habitat structure and quality in determining species distribution within human-dominated landscapes.

The relationship between species abundance and habitat selection is complex and often mediated by both biotic and abiotic factors that influence resource availability and environmental suitability (Felton *et al*., 2021). In heterogeneous landscapes, such as those combining human settlements and farmlands, birds must navigate a mosaic of habitat types that differ in structure, resource distribution, and levels of disturbance (Morelli *et al*., 2021; Aronson *et al*., 2020). As a result, patterns of abundance are not random but reflect the outcomes of habitat selection processes, where individuals preferentially occupy areas that offer optimal conditions for survival and reproduction ((Felton *et al*., 2021). This is particularly relevant for small passerines like the Red-cheeked Cordon-bleu, whose abundance is closely linked to vegetation structure and food availability. In human-modified landscapes, habitat selection is often influenced by anthropogenic factors that alter the natural distribution of resources and environmental conditions (Benítez-López *et al*., 2021; Samia *et al*., 2022). For instance, urbanization can lead to the loss of natural habitats and the creation of artificial environments that may either support or exclude certain bird species depending on their ecological traits (Aronson *et al*., 2020; Callaghan *et al*., 2024). Similarly, agricultural activities can modify vegetation structure and resource availability, thereby influencing habitat suitability for different species (Morelli *et al*., 2021; Tryjanowski *et al*., 2020). Birds that are able to adapt to these changes by exploiting new resources or tolerating disturbance are more likely to maintain or increase their abundance, while less adaptable species may decline.

Measuring species abundance and habitat selection in ecological studies typically involves standardized sampling methods such as point counts and transect surveys, which allow researchers to estimate population size and distribution across different habitat types (Sutherland *et al*., 2020; Johnston *et al*., 2022). These methods are often combined with habitat assessments to quantify environmental variables and analyze their relationship with species abundance (Felton *et al*., 2021). The point count method involves recording all individuals of a target species seen or heard from a fixed location within a specified time period, allowing for standardized comparisons across different sites (Callaghan *et al*., 2024; Morelli *et al*., 2021).

This method is particularly useful in heterogeneous landscapes such as rural settlements and farmlands, where habitat patches may be small and spatially discontinuous (Staley *et al*., 2023; Aronson *et al*., 2020). The line transect method, on the other hand, involves systematic movement along predetermined paths while recording bird observations at varying distances from the transect line (Johnston *et al*., 2022; Sutherland *et al*., 2020). This approach is effective for covering larger spatial areas and capturing variation in bird distribution across habitat gradients (Morelli *et al*., 2021; IPBES, 2021). Both methods require careful consideration of detectability, as factors such as vegetation density, observer experience, and environmental conditions can influence the probability of detecting individuals (Callaghan *et al*., 2024; Felton *et al*., 2021). Failure to account for detectability can lead to biased abundance estimates, particularly in comparative studies between structurally different habitats (Redhead *et al*., 2021; Staley *et al*., 2023). In the context of the Red-cheeked Cordon-bleu, species abundance is likely to be influenced by the availability of suitable foraging grounds, nesting sites, and levels of disturbance within different habitat types (Craig *et al*., 2020; Evans *et al*., 2022). Human settlements and farmlands may differ significantly in these attributes, leading to variations in habitat selection and, consequently, differences in abundance between these environments (Morelli *et al*., 2021). Statistical approaches such as regression analysis and occupancy modeling are then used to identify key predictors of abundance and to evaluate the strength of habitat associations (Callaghan *et al*., 2024). Such methodologies are essential for generating robust ecological data and for testing hypotheses related to habitat selection

## MATERIALS AND METHODS

### Study Area

The study was carried out in Laminga Village, a rural settlement located within Jos-East Local Government Area of Plateau State, North-Central Nigeria, and it forms part of the Guinea savanna ecological zone characterized by a mixture of grassland, scattered shrubs, and anthropogenically modified landscapes (FAO, 2021; World Bank, 2023). The study area lies within a transition zone between more densely vegetated upland environments and human-dominated agricultural landscapes, making it ecologically suitable for examining species responses to varying degrees of habitat modification (IPBES, 2021; Shackleton *et al*., 2021). The ecological significance of the study area is further enhanced by its position within a savanna ecosystem that supports a diverse assemblage of bird species adapted to open and semi-open habitats (BirdLife International, 2023; Evans *et al*., 2022). As shown in Figure 1, the region is typified by a mosaic of human settlements, small-scale farmlands, and remnant natural vegetation patches, which collectively provide heterogeneous habitats for avian species such as the Red-cheeked Cordon-bleu (*Uraeginthus bengalus*) (BirdLife International, 2023; Morelli *et al*., 2021).

**Figure 1:**
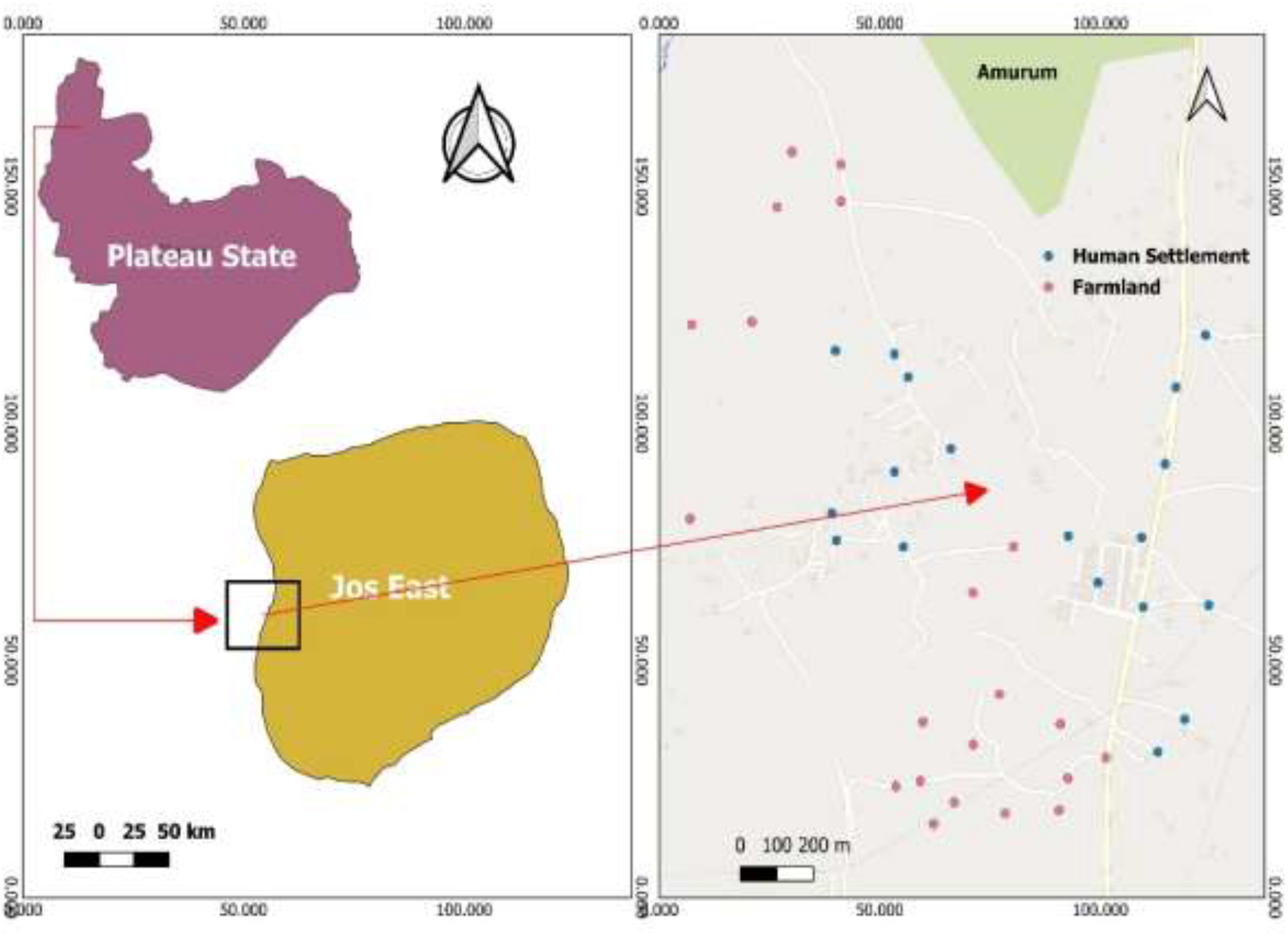
Map of the study site showing the sampled locations (Generated using QGIS version 3.40.1-Bratislava)

**Figure 2:**
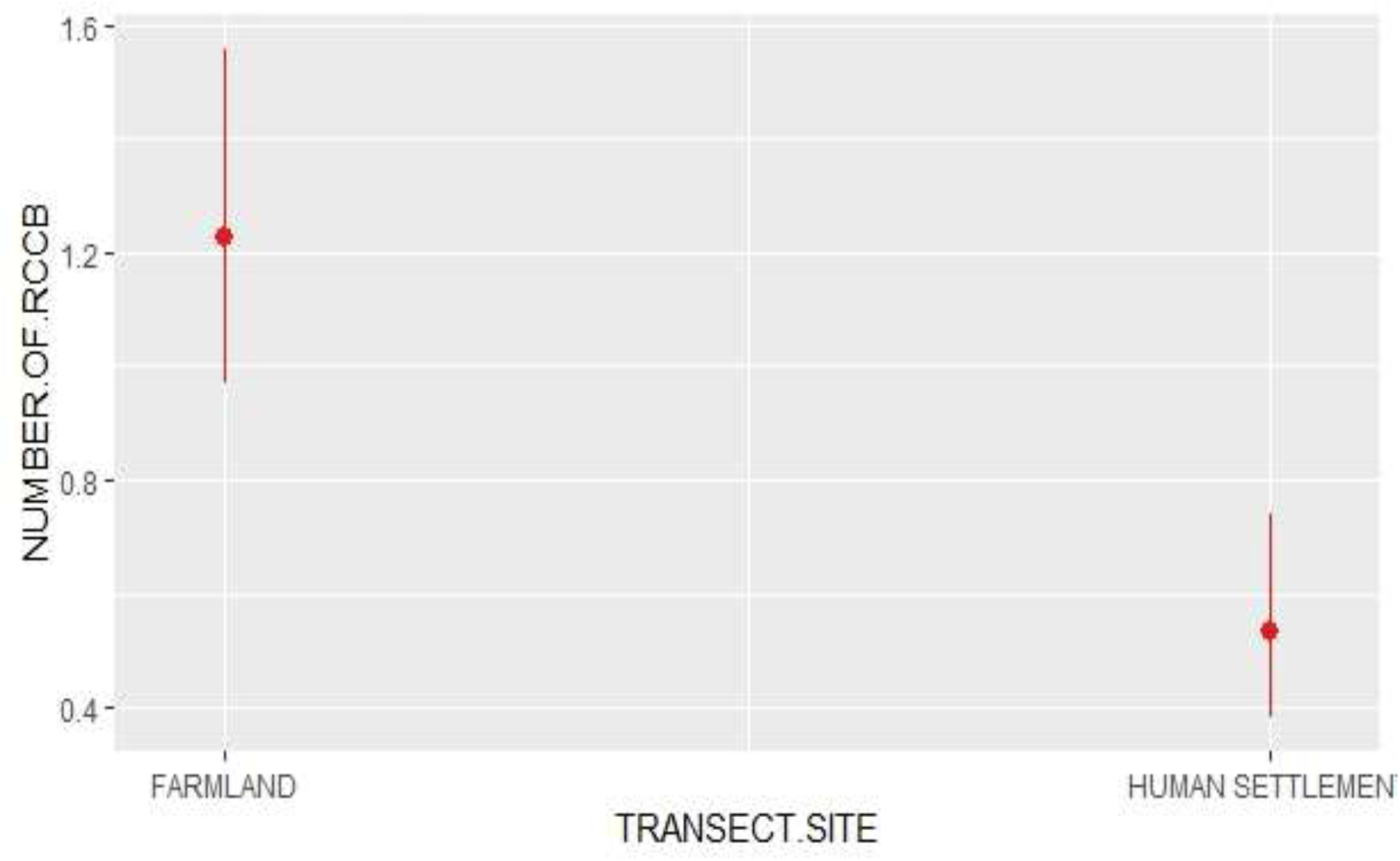
Comparison of abundance of Red-cheeked Cordon-bleu in Human settlements and Farmlands.

### Study Duration

The study was conducted over a period of three weeks in November 2024, a period selected to coincide with stable post-rainy season conditions when vegetation structure is relatively established and bird detectability is high (Evans *et al*., 2022; Morelli *et al*., 2021).

### Bird Sampling Techniques

Data collection was carried out during peak avian activity periods, specifically early morning hours between 6:30 am and 9:30 am, and late afternoon hours between 4:00 pm and 6:00 pm, to maximize detection probability and reduce temporal bias in abundance estimates. A total of 21 transects were established across the study area, each measuring approximately 200 meters in length and subdivided into 50-meter segments to allow for systematic sampling of bird abundance and habitat variables. Within human settlement areas, 10 transects of 200 meters each were established, while farmland areas comprised 8 transects of 200 meters, 2 transects of 150 meters, and 1 transect of 100 meters, all positioned at least 200 meters away from settlement boundaries to minimize spatial overlap effects between habitat types. This design ensured adequate representation of both habitat categories for comparative analysis (Morelli *et al*., 2021; IPBES, 2021). Bird abundance in this study was quantified as the number of individual Red-cheeked Cordon-bleus (*Uraeginthus bengalus*) detected per sampling unit within each transect segment, following standard practices in avian ecological surveys where count data are used as proxies for relative population density (Sutherland *et al*., 2020; Johnston *et al*., 2022).

### Data Analysis

The generated data was cleaned in MS Excel, and imported into R version 4.5.1 statiatical package for analysis. Statistical analysis was conducted using Generalized Linear Models (GLM) with a Poisson distribution and log-link function to examine the relationship between Red-cheeked Cordon-bleu abundance and explanatory variables. A model was fitted to assess the effect of landscape type and transect length on abundance.

## RESULTS

### Abundance of Red-cheeked Cordon-bleu in Settlements and Farmlands

The comparative analysis of Red-cheeked Cordon-bleu (*Uraeginthus bengalus*) abundance between human settlements and farmlands varies demonstrating a statistically significant difference in species distribution across the two habitat types. The predicted counts derived from the Poisson Generalized Linear Model (GLM) revealed a clear spatial pattern in species distribution, indicating a higher abundance in farmland habitats compared to human settlements. As shown in Figure 1, the mean predicted abundance of the species in farmlands was notably higher, with values approximating 1.2 individuals per sampling unit, whereas human settlements exhibited a substantially lower mean abundance, approximately 0.5 individuals per sampling unit. This difference highlights a strong habitat preference or ecological suitability associated with farmland environments. Statistically, this difference was supported by the regression coefficient (β = −0.84), with a 95% confidence interval ranging from −1.22 to −0.47 and a highly significant p-value (p < 0.001) as shown in Table 1. The negative coefficient indicates that the reference category (human settlements) had a significantly lower abundance relative to farmlands. Since the confidence interval does not cross zero and the p-value is well below the conventional significance threshold (0.05), this result confirms that habitat type is a strong predictor of Red-cheeked Cordon-bleu abundance.

**Table 1:** Effects of Habitat Type on the Abundance of Red-cheeked Cordon-bleu.

| **Predictor** | **$\beta$ (Estimate)** | **SE** | **z-value** | **p-value** |
| --- | --- | --- | --- | --- |
| Intercept | -5.075 | 1.515 | -3.350 | <0.001 |
| Human settlement | -0.836 | 0.192 | -4.359 | <0.001 |
| Transect length | 0.028 | 0.008 | 3.600 | <0.001 |
| Model statistics: Null deviance = 159.88, Residual deviance = 125.85, AIC = 306.32 |  |  |  |  |

## DISCUSSION

The results of this study demonstrate a clear and statistically significant difference in the abundance of the Red-cheeked Cordon-bleu (*Uraeginthus bengalus*) between human settlements and farmlands, with farmlands supporting substantially higher abundance. This pattern reflects fundamental ecological principles relating to habitat suitability, resource availability, and disturbance gradients in human-modified landscapes. The higher abundance observed in farmlands can be attributed primarily to greater availability of food resources and more favorable habitat structure. As a granivorous species, the Red-cheeked Cordon-bleu depends heavily on seeds from grasses and crops, which are more abundant and accessible in agricultural environments. Farmlands also provide heterogeneous vegetation structure, including grasses, shrubs, and crop residues, which enhance foraging efficiency and offer protection from predators (Morelli *et al*., 2021; Evans *et al*., 2022). In contrast, human settlements are characterized by reduced vegetation cover, increased habitat fragmentation, and higher levels of anthropogenic disturbance. These factors collectively reduce habitat suitability for many bird species, particularly those that rely on ground vegetation for feeding and shelter. The significantly lower abundance observed in settlements aligns with findings that urbanization tends to favor a limited number of generalist species while reducing the abundance of others (Callaghan *et al*., 2024; Aronson *et al*., 2020).

The negative regression coefficient associated with settlements further confirms that habitat type is a strong determinant of abundance. This suggests that even though the species is capable of persisting in human-modified environments, its optimal ecological niche remains within less disturbed, vegetation-rich habitats. Such patterns are consistent with broader ecological theory, which indicates that species abundance declines along gradients of increasing urbanization due to reduced resource availability and increased disturbance (IPBES, 2021; Evans *et al*., 2022). Additionally, the spatial separation of farmland transects from settlements likely reduced edge effects and direct human interference, further enhancing habitat quality. This supports the concept that landscape configuration, not just habitat type, plays a role in shaping species distribution (Morelli *et al*., 2021).

## Conclusion

Habitat type plays a dominant role in determining the abundance of the Red-cheeked Cordon-bleu. The species was significantly more abundant in farmlands than in human settlements, indicating a clear preference for environments that maintain greater vegetation cover and lower levels of continuous disturbance. This finding aligns with ecological theory suggesting that agricultural landscapes, particularly those with heterogeneous vegetation, can support higher bird abundance compared to more intensively modified settlement areas. The results therefore confirm that farmlands within the study area serve as more suitable habitats for the species.

